# Fine-tuned adaptation of embryo-endometrium pairs at implantation revealed by gene regulatory networks Tailored conceptus-maternal communication at implantation

**DOI:** 10.1101/427310

**Authors:** Fernando H. Biase, Isabelle Hue, Sarah E. Dickinson, Florence Jaffrezic, Denis Laloe, Harris Lewin, Olivier Sandra

**Affiliations:** Department of Animal Sciences, Auburn University, Auburn, AL, USA 36839; INRA, UMR 1198 Biologie du Développement et Reproduction, Ecole Nationale Vétérinaire d’Alfort, F-78350 Jouy-en-Josas, France 78352; UMR Génétique Animale et Biologie Intégrative, Institut National de la Recherche Agronomique, Jouy en Josas, France 78352; Department of Evolution and Ecology, University of California, Davis, CA, USA 95616

**Keywords:** early pregnancy, system biology, embryo, cattle

## Abstract

Interactions between embryo and endometrium at implantation are critical for the progression and the issue of pregnancy. These reciprocal actions involve exchange of paracrine signals that govern implantation and placentation. However, it remains unknown how these interactions between the conceptus and the endometrium are coordinated at the level of an individual pregnancy. Under the hypothesis that gene expression of endometrium is dependent on gene expression of extraembryonic tissues, we performed an integrative analysis of transcriptome profiles of paired conceptuses and endometria obtained from pregnancies initiated by artificial insemination. We quantified strong dependence (|r|>0.95, eFDR<0.01) in transcript abundance of genes expressed in the extraembryonic tissues and genes expressed in the endometrium. The profiles of connectivity revealed distinct co-expression patterns of extraembryonic tissues with caruncular and intercaruncular areas of the endometrium. Notably, a subset of highly co-expressed genes between conceptus (n=229) and caruncular areas of the endometrium (n=218, r>0.9999, eFDR<0.001) revealed a blueprint of gene expression specific to each pregnancy. Functional analyses of genes co-expressed between conceptus and endometrium revealed significantly enriched functional modules with critical contribution for implantation and placentation, including “in utero embryonic development”, “placenta development” and “regulation of transcription”. Functional modules were remarkably specific to caruncular or intercaruncular areas of the endometrium. The quantitative and functional association between genes expressed in conceptus and endometrium emphasize a coordinated communication between these two entities in mammals. To our knowledge, we provide first evidence that implantation in mammalian pregnancy relies on the ability of the conceptus and the endometrium to develop a fine-tuned adaptive response characteristic of each pregnancy.

## INTRODUCTION

In mammals, pregnancy recognition requires a tightly synchronized exchange of signals between the competent embryo and the receptive endometrium. The initiation of this signaling is triggered by key factors produced by the conceptus (1, 2) which are translated by the endometrial cells into actions that will condition the trajectory of embryo development as well as progeny phenotype. In mammalian species, including human, rodents and ruminants, the delicate balance in embryo-maternal communication is affected by the way the embryos are generated (natural mating, artificial insemination, *in vitro* fertilization somatic cell nuclear transfer) and by the sensor-driver properties of the endometrium defined by intrinsic maternal factors (i.e.: maternal metabolism, ageing) and environmental perturbations (i.e.: pathogens, nutrition) (3-5). The concept of sensor property applied to the mammalian endometrium was first proposed in a pioneer paper as was suggested the notion of endometrial plasticity (6). This property was recently confirmed *in vitro* with an aberrant responsiveness of human endometrial stromal cultured cells in the context of recurrent pregnancy loss (7). Nevertheless, it remains unaddressed whether the mammalian endometrium is able to develop an adaptive embryo-tailored response in a normal pregnancy.

In mammalian reproduction, sheep and cattle are research models that have relevantly contributed key insights in the understanding of molecular and physiological pregnancy-associated mechanisms, including the deciphering of embryo-endometrium interactions (8, 9). In the bovine species, by gestation days 7-8, the blastocyst enters the uterine lumen. After hatching by days 8-9, the outer monolayer of trophectoderm cells establishes direct contact with the luminal epithelium of the endometrium (10). On gestation days 12-13, the blastocyst is ovoid in shape (~2-5 mm) and transitions into a tubular shape by days 14-15. By day ~15, rapidly proliferating trophectoderm cells of the extra-embryonic tissues synthesize and release IFNT (11-15), which is the major pregnancy recognition signal in ruminants (1, 9, 16, 17). The disrupted release of the oxytocin-dependent pulses of prostaglandin F2 alpha (18) allows maintenance of progesterone production by a functional corpus luteum (18), which is critical for the establishment and progression of pregnancy (1, 4, 9, 12, 14, 15, 19). IFNT actions include induction of numerous classical and non-classical IFN-stimulated genes and stimulation of progesterone-induced genes that encode proteins involved in conceptus elongation and implantation (4). IFNT-regulated genes have diverse actions in the endometrium that are essential for conceptus survival and pregnancy establishment (12). Other paracrine signals such as prostaglandins and cortisol have regulatory effects on conceptus elongation and endometrium remodeling (20). More recently, the identification of potential ligand-receptor interactions between the conceptus and endometrium (21) and the secretion of proteins and RNAs through exosomes (22, 23) have expanded the field of possibilities by which the conceptus and endometrium interact prior to and during implantation.

The cross-talk between the conceptus and the endometrium is associated with the expression and regulation of a wealth of genes in each entity (24, 25). The nature of the conceptus modifies gene expression of the endometrium in cattle (6, 26, 27) and decidualizing human endometrial stromal cells (28). Similarly, the endometrium from dams with different fertility potentials (29) or metabolic status (30) influences the gene expression of the conceptus. Despite the growing evidence of the interactions between conceptus and endometrium at the level of gene regulation, the pathways and the functions that result from this interaction have yet to be unveiled. Furthermore, the lack of integrated analysis between paired conceptus and endometrium has made it challenging to advance our understanding of the functional interactions between these two entities in normal pregnancies.

Here, we hypothesized that gene expression of extraembryonic tissue is not independent from gene expression of endometrium. In the present study, we carried out an integrative analysis of transcriptome profiles of paired conceptuses and endometria at the onset of implantation aiming at the identification of regulatory pathways that have coordinated expression between the conceptus and endometrium in normal pregnancies. Surprisingly, our results show that at gestation day 18 in cattle, several hundred genes have an expression profile in conceptus and caruncular areas of the endometrium that is unique to each pregnancy. Analyses of genes co-expressed between the conceptus and the paired-associated endometrium revealed significantly enriched functional modules with critical contribution for implantation and placentation. Our data provide evidence that successful implantation in mammalian pregnancy relies on the ability of the endometrium to elicit a fine-tuned adaptive response to the conceptus.

## RESULTS

### Data overview

We analyzed the RNA-sequencing data that consisted of samples collected from five cattle pregnancies terminated at gestation day 18 (GSE74152 (26)). The conceptus was dissected, and transcriptome data was generated for extraembryonic tissue; whereas the endometrium was dissected into caruncular (gland-free) and intercaruncular (containing endometrial glands) areas, and transcriptome data was generated from both regions of the endometrium (Fig 1A). Therefore, the dataset analyzed was comprised of three samples collected from each pregnancy: extraembryonic, caruncular, and intercaruncular tissues (Fig 1B). Alignment of the sequences to the *Bos taurus* genome (UMD 3.1) resulted into an average of 22, 31.4, and 34.6 million uniquely mapped reads for extraembryonic (n = 5), caruncular (n = 5), and intercaruncular (n = 5) tissues, respectively. We quantified the transcript abundance of 9548, 13047, and 13051 genes in extraembryonic, caruncular, and intercaruncular tissues, respectively (Fig 1C). Unsupervised clustering of the samples based on their transcriptome data separated the samples obtained from the conceptus from the endometrial samples and further distinguished caruncular from intercaruncular endometrial samples (Fig 1D).

**Fig 1.**
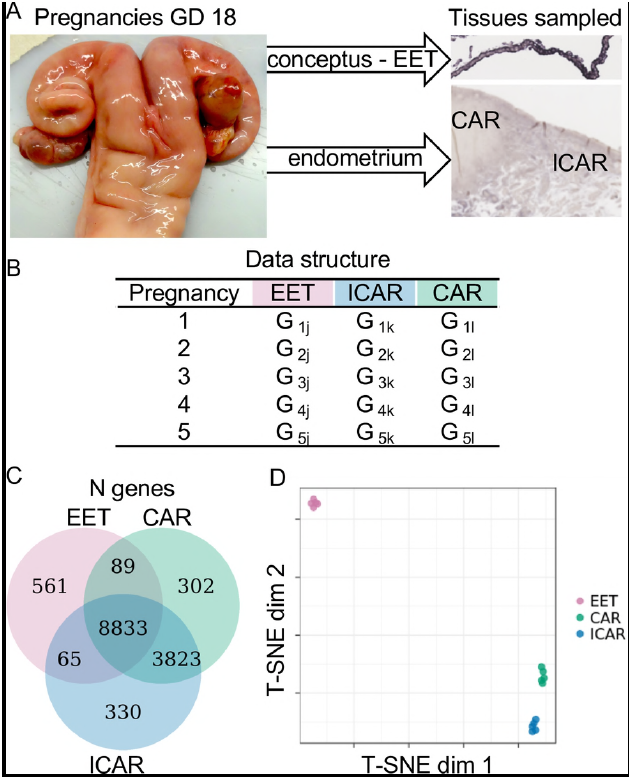
Transcriptome profiling of conceptus and endometrium collected from gestation day 18. (A) Representative images of pregnant uterus and micrograph identifying the tissues from which RNA-seq data were used in this study. (B) Data structure used in this study. Data on genome-wide transcript abundance was obtained from extraembryonic tissue (EET) and endometrium (caruncular (CAR), intercaruncular (ICAR) tissues) from five pregnant uteri. (C) Number of genes with transcript abundance quantified in each sample. (D) Dimensionality reduction of the RNA-seq data for special visualization of the sample distribution.

### Correlated gene expression between conceptus and endometrium

The associated expression between two genes can be assessed by correlative metrics (31) within (32, 33) or between tissues (33, 34). Thus, we calculated Pearson’s coefficient of correlation (r (35)) to test whether there is association between the transcript abundance of genes expressed in extraembryonic tissue and endometrium (caruncular or intercaruncular tissues). We reasoned that under a null hypothesis, the abundance of a gene expressed in extraembryonic tissue (G_j_) would have no association with the abundance of a gene expressed in endometrium (G_k_, or G_l_), for example: 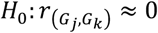. On the other hand, under the alternative hypothesis 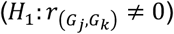, two genes display co-expression (35).

The distribution of correlation coefficients for all pairs of genes expressed in extraembryonic and caruncular tissues averaged 0.13 (Fig 2A), and the equivalent distribution obtained for all pairs of genes expressed in extraembryonic and intercaruncular tissues averaged 0.03 (Fig 2B). Both distributions deviated significantly from a distribution obtained from shuffled data that disrupted the pairing of the conceptus and endometrium (P < 2.2^-16^, S1 Fig). We calculated the empirical FDR (eFDR) and noted that absolute correlation coefficients in both distributions were highly significant when greater than 0.95 (eFDR < 0.007, S2 Fig, S1 Table). Of note, S3 Fig and S4 Fig present examples of pairs of genes we identified with the highest positive and negative correlation coefficients, which fit the alternative hypothesis 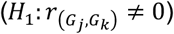 and examples of pairs of genes that show correlation coefficients close to zero fitting the null hypothesis 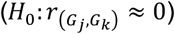.

**Fig 2.**
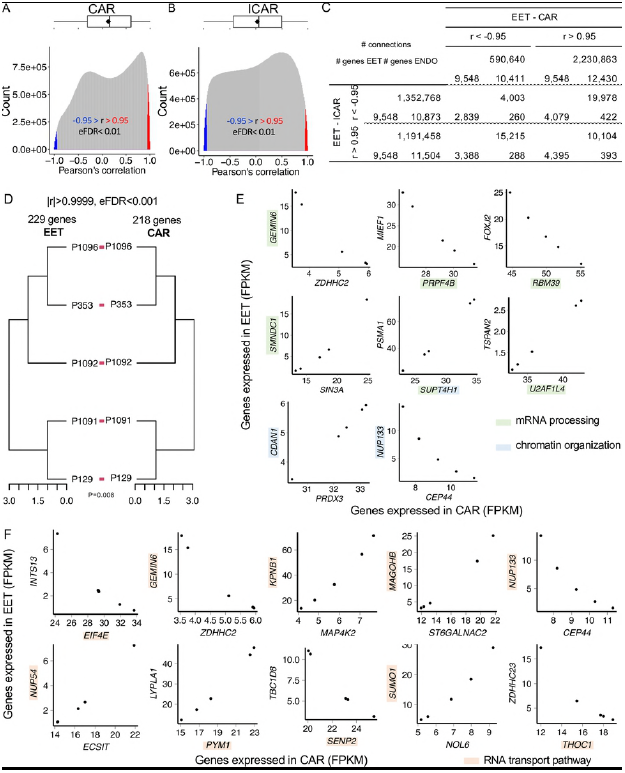
Co-expression analysis between extraembryonic tissue and endometrium. Distribution of the correlation coefficients for genes expressed in extraembryonic (EET) and caruncular (CAR) tissues (A) or intercaruncular (ICAR) tissues (B). (C) Number of genes expressed in either EET, CAR or ICAR that participate in significant correlation connections involving conceptus and endometrium. (D) Tanglengram of EET and CAR tissues formed by genes with strong co-expression. (E) Scatterplot of pairs of genes expressed in EET and CAR with at least one gene involved in “mRNA processing” or “chromatin organization”. (F) Scatterplot of pairs of genes expressed in EET and CAR and highly correlated with at least one gene involved in “RNA transport pathway”.

Notably, all 9548 genes expressed in extraembryonic tissue where positively (r > 0.95) and negatively (-0.95 > r) correlated with genes expressed in caruncular or intercaruncular tissues (Fig 1C). Eighty percent and 95% of the genes expressed in caruncle tissues were negatively and positively correlated with genes expressed in extraembryonic tissue. Similarly, 83% and 88% of the genes expressed in intercaruncular tissues were negatively and positively correlated with genes expressed in extraembryonic tissue (Fig 1C). The distribution of degrees of connectivity for significant correlations (|r| > 0.95, eFDR < 0.01) between extraembryonic and caruncular tissues was not equivalent to the distribution observed between extraembryonic and intercaruncular tissues (P < 2.2^-16^). The genes expressed in extraembryonic tissue were significantly correlated with 295 genes expressed in caruncular tissues on average (median = 101). Eleven genes were significantly correlated with over 2300 genes in caruncular tissues (i.e. *AREG, EGR1, PEX3, GAN*, S5A Fig). The genes expressed in extraembryonic tissue were significantly correlated with 266 genes expressed in intercaruncular tissues on average (median = 252). Eight genes were significantly correlated with over 750 genes in intercaruncular tissues (i.e.: *WNT5B, WNT7B, ROR2, DPEP1, GJB3*, S4B Fig). These results strongly suggest different patterns of gene co-expression between extraembryonic and caruncular or intercaruncular tissues.

We then examined if genes co-expressed in extraembryonic tissue and endometrium have expression patterns that are unique to pregnancies. We identified 229 and 218 genes expressed in extraembryonic and caruncular tissues, respectively (|r| > 0.9999, eFDR < 0.001, S1 Table), whose expression profiles produced equivalent dendrograms for extraembryonic and caruncular tissues independently (P = 0.008, Fig 2D). Functional investigation of these 441 genes identified significant enrichment in the biological processes “mRNA processing” (*GEMIN6, PRPF4B, RBM39, SMNDC1, SUPT4H1, U2AF1L4*, FDR = 0.13, Fig 2E), “chromatin organization” (*CDAN1, NUP133, SUPT4H1*, FDR = 0.13, Fig 2E), and “protein autoubiquitination” (*CNOT4, MARCH5, UHRF1*). We also interrogated the KEGG pathways database and identified an enrichment for the “RNA transport” pathway (*EIF4E, GEMIN6, KPNB1, MAGOHB, NUP133, NUP54, PYM1, SENP2, SUMO1, THOC1*, FDR = 0.06, Fig 2F). We did not identify groups of genes co-expressed in extraembryonic and intercaruncular tissues capable of producing dendrograms that mirrored each other. These results demonstrate that genes highly co-expressed between extraembryonic and caruncular tissues form a signature that independently distinguishes pregnancies in an equivalent manner.

### Visualization of co-expressed networks in extraembryonic tissue and endometrium

Our analysis was not an exhaustive evaluation of all potential co-expression networks that exist between conceptus and endometrium. Thus, we developed a web interface for dynamic and interactive data visualization based on the co-expression analysis conducted in the present study (36, 37) (https://biaselab.shinyapps.io/eet_endo/). The public access to this web application allows a user to produce networks for genes of their choosing. Furthermore, each network is accompanied by supporting data such as scatter plots and heatmaps of the gene expression values. The raw data and codes for reproduction of this interface can be downloaded from a GitHub repository (https://github.com/BiaseLab/eet_endo_gene_interaction).

### Functional networks between extraembryonic and caruncular tissues

We investigated the transcriptome-wide interactions between extraembryonic and caruncular or intercaruncular tissues independently. The clustering of genes based on co-expression is a powerful means to understand coordinated gene functions (38), thus we used the matrix with correlation coefficients to cluster extraembryonic, caruncular, and intercaruncular tissues independently.

The heatmap resultant of clustering the two datasets (extraembryonic and caruncular tissues) showed the formation of an organized co-expression network between the genes expressed in extraembryonic and caruncular tissues (Fig 3A). We identified 36 clusters formed by the genes expressed in extraembryonic tissue that presented enrichment for several biological processes (FDR < 0.2, Fig 3B), where we identified several genes expressed in extraembryonic tissue significantly co-expressed with genes expressed in caruncular tissues (see S1 Data for a list of genes). For instance, 142 genes associated with regulation of transcription were identified across clusters 1, 12, 30, 38, and 54. Eighty-two genes were associated with signal transduction across clusters 1, 21, 27, and 71. Interestingly, 26 genes associated with “in utero embryonic development” were identified in cluster 1.

**Fig 3.**
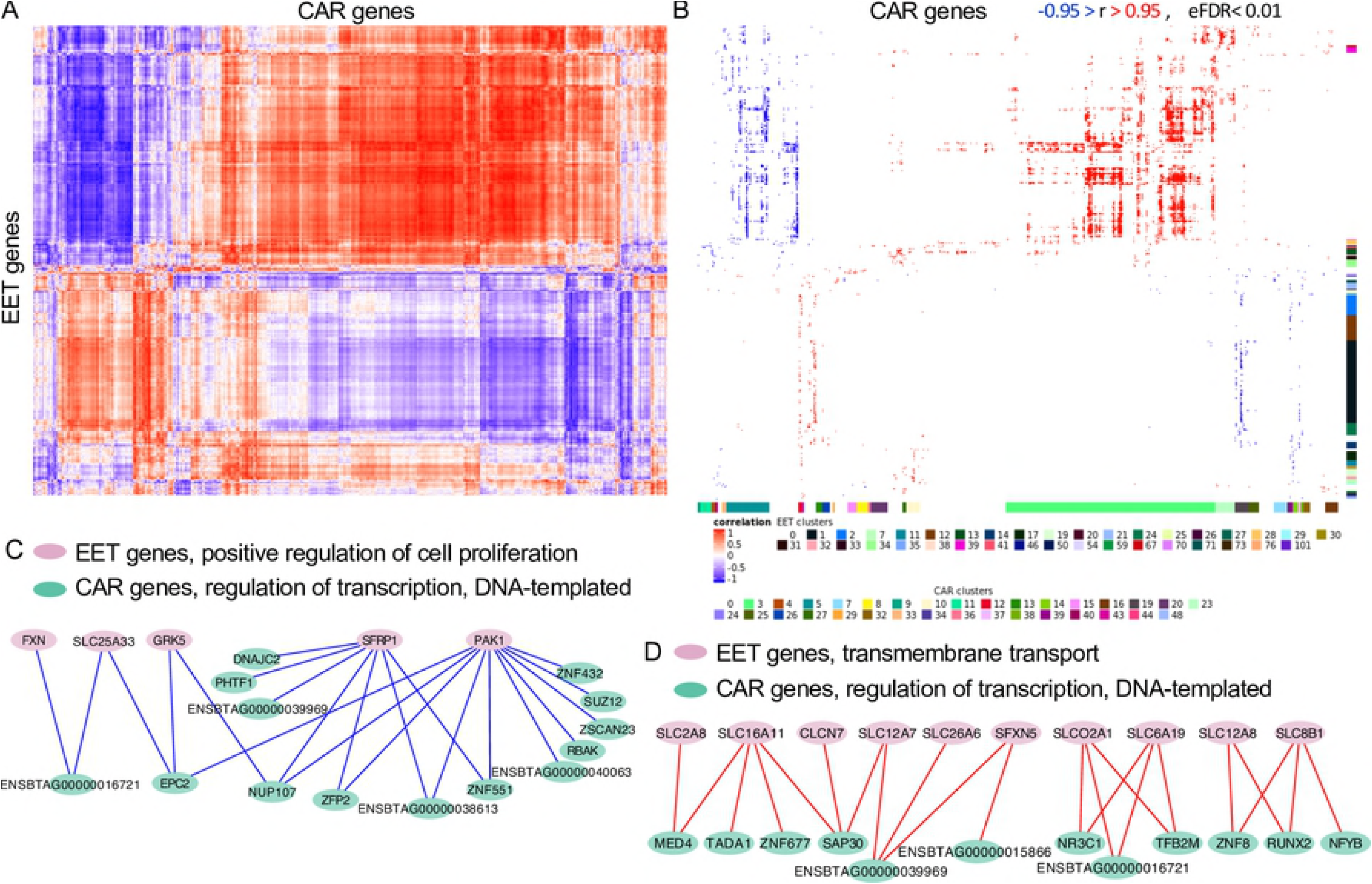
Functional analysis of co-expressed genes between extraembryonic (EET) and caruncular (CAR) tissues. (A) Heatmap produced by the correlation coefficients and independent clustering of EET and CAR. (B) Gene ontology analysis of the cluster formed. Only significant coefficients of correlation are shown (|r| > 0.95, eFDR<0.01). The colored bar on the right of the heatmap indicates clusters of genes expressed in EET for which biological processes were significant. The colored bar on bottom of the heatmap indicates clusters of genes expressed in caruncle for which biological processes were significant. The colored squares at the bottom of the image identify the cluster number with the color observed on the bars. See S1 Data and S2 Data for details on the cluster identification, biological processes and genes. (C,D) Model of functional co-expression networks possibly formed between EET and CAR.

The clustering of genes expressed in caruncular tissues according to their co-expression with extraembryonic tissue genes resulted in the identification of 32 clusters presenting enrichment (FDR<0.2) for several Biological processes (Fig 3A, S2 Data). Among the genes forming significant co-expression with extraembryonic tissue, we identified 96 genes in cluster 3 associated with “intracellular protein transport”, as well as 111 and four genes associated with regulation of transcription in clusters 4 and 5, respectively. Notably, ten genes on cluster 15 were associated with “defense response to virus”, and the annotated genes are known to be stimulated by interferon-tau (*IFIT1, IFIT3, IFIT5, ISG15, MX1, MX2, OAS1Y, RSAD2*, S2 Data).

Next, we intersected the results of gene ontology enrichment obtained from clustering extraembryonic and caruncular tissues. We identified several biological processes on both datasets with co-expressing genes expressed in extraembryonic and caruncular tissues (S3 Data). Based on the number of genes and direction of connections, two pairs of biological processes are noteworthy. First, five genes associated with “positive regulation of cell proliferation” in extraembryonic tissue form negative co-expression connections 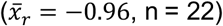 with 14 genes associated with “regulation of transcription, DNA-templated” expressed in caruncle (Fig 3C). Second, ten genes associated with “transmembrane transport” in extraembryonic tissue form positive co-expression connections 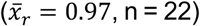 with 12 genes associated with “regulation of transcription, DNA-templated” expressed in caruncle (Fig 3D). These results are coherent with a co-expression between genes expressed in extraembryonic and caruncular tissues, with functional implications to conceptus attachment and implantation.

### Functional networks between extraembryonic and intercaruncular tissues

The independent clustering of the correlation coefficients obtained from the genes expressed in extraembryonic and intercaruncular tissues also evidenced an organized co-expression network between the two tissues (Fig 4A). Twelve clusters formed by genes expressed in the extraembryonic tissue presented enrichment for biological processes (FDR < 0.2, Fig 4B, see S4 Data for a list of genes). Interestingly, there were 85 and 27 genes associated with “mRNA processing” and “stem cell population maintenance”, respectively on cluster 3. On cluster five, we identified 12 genes associated with “negative regulation of cell proliferation” and seven genes associated with “regulation of receptor activity”. On cluster eight, 5 genes were associated with “placenta development” (*ADA, CCNF, DLX3, PHLDA2, RXRA*). On cluster 17, eight genes were associated with “regulation of transcription, DNA-templated”.

**Fig 4.**
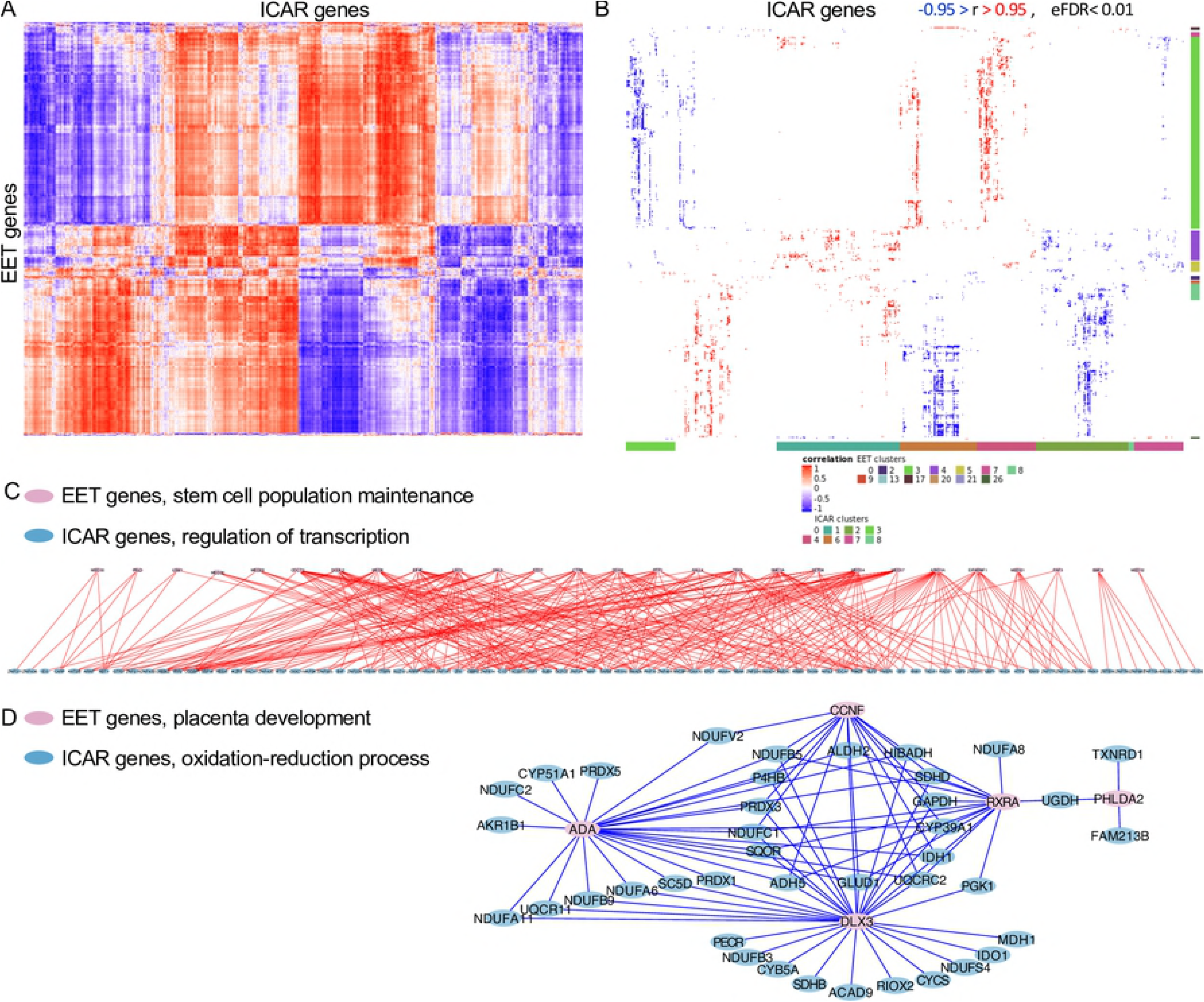
Functional analysis of co-expressed genes between extraembryonic (EET) and intercaruncular (ICAR) tissues. (A) Heatmap produced by the correlation coefficients and independent clustering of EET and ICAR. (b) Gene ontology analysis of the cluster formed. Only significant coefficients of correlation are shown (|r| > 0.95, eFDR<0.01). The colored bar on the right of the heatmap indicates clusters of genes expressed in EET for which biological processes were significant. The colored bar on bottom of the heatmap indicates clusters of genes expressed in intercaruncle for which biological processes were significant. The colored squares at the bottom of the image identify the cluster number with the color observed on the bars. See S4 Data and S5 Data for details on the cluster identification, biological processes and genes. (C,D) Model of functional co-expression networks possibly formed between EET and ICAR. See S6 fig for an enlarged version of panel C.

The clusters formed by intercaruncular genes co-expressed with extraembryonic tissue genes also highlighted significant enrichment of biological processes (FDR < 0.2, Fig 4A, see S5 Data for a list of genes). For instance, clusters one and six contained 145 and 63 genes associated with regulation of transcription, respectively. Interestingly, on cluster two, there were 149, 23, 22, and 16 genes associated with “oxidation-reduction process”, “cell redox homeostasis”, “electron transport chain”, and “tricarboxylic acid cycle”. Cluster four contained 63 genes associated with “regulation of transcription”, and cluster seven contained 11 genes associated with “fatty acid beta-oxidation”.

The intersection of the genes identified in enriched biological processes in clusters formed by extraembryonic and intercaruncular tissues revealed several potential functional co-expression networks between these two tissues (S6 Data). Notably, several of the intersecting categories involved processes associated with regulation of transcription or oxidation-reduction on the intercaruncular side. For instance, 28 genes associated with “stem cell population maintenance” and expressed in extraembryonic tissue presented positive co-expression 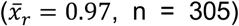 with 83 genes associated with “regulation of transcription” and expressed in intercaruncular tissues (Fig 4D). Five genes associated with “placenta development” and expressed in extraembryonic tissue presented negative co-expression 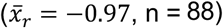 with 41 genes associated with “oxidation-reduction process” and expressed in intercaruncular tissues (Fig 4E).

## DISCUSSION

In mammals and particularly in the bovine species, a large body of gene expression data was produced at various steps of early pregnancy derived from *in vitro* or *in vivo* produced embryos (6, 26, 27, 39), varied physiological status of the dam (40), and fertility classified heifers (29). Altogether, results based on groups analyses (conceptus or endometrium) have demonstrated different degrees of interactions between the conceptus and endometrium at the initial phases of implantation. In the present study, our objective was to shed light on the subtle interactions between the extraembryonic tissue of a conceptus and the endometrial tissue of the uterus hosting this conceptus in normal pregnancy using paired co-expression analyses of gene transcript abundances. Our analyses were carried out using biological material collected from the single conceptus and the endometrium from the same pregnancy, a critical aspect to determine the cross-talk during implantation at the level of one individual pregnant female.

Our analyses of transcriptome data from conceptus and endometrium pairs identified key signatures of gene expression that are likely to be linked to the success of pregnancy recognition and implantation. A large proportion of all genes quantified in extraembryonic tissue and endometrium have transcript abundances that were not independent. Furthermore, the dependency observed for the abundance of transcripts between extraembryonic tissue and endometrium varied with morphologically and physiologically distinct areas of the endometrium, namely caruncular and intercaruncular tissues. For instance, there were twice as many highly positive (r > 0.95) and approximately half the number of highly negative (r < -0.95) co-expressing connections between extraembryonic and caruncular tissues compared to extraembryonic and intercaruncular tissues. These results greatly expand previous findings that the conceptus triggers distinct molecular responses in caruncular and intercaruncular tissues (6, 26, 41, 42).

During the elongation phase, the mural trophoblast proliferates rapidly (12, 25, 43) while maintaining its pluripotency (44). This period of development is modulated by dynamic regulation of gene expression (43) whereby metabolically active trophoblastic cells (45, 46) rely on the uptake of nutrients from the uterine luminal fluid (47). Our results show that caruncular and intercaruncular tissues have an active role in the programing of those functions, as several genes related with gene regulation, signal transduction, cellular proliferation, maintenance of stem cell population, and transmembrane transport are also co-expressed with genes expressed in the endometrium. The importance of gene co-regulation between extraembryonic tissue and endometrium was further supported by the identification of 26 genes associated with “in utero embryonic development” and five genes associated with “placenta development” co-regulated with genes expressed in caruncle and intercaruncle, respectively.

Among the genes expressed in caruncular or intercaruncular tissues that were co-expressed with extraembryonic tissues, it was noticeable that several genes were associated with regulation of gene expression. This finding is in line with former publications reporting that the regulatory network needed for endometrial remodeling (48) during attachment is conceptus-dependent. In the caruncular tissue, we specifically identified 15 genes associated with “defense response to virus”, of which eight genes had their expression modulated by interferon-tau, produced by the trophoblast between gestation days 9 and 25 (49). This result provide additional knowledge on the biological actions of interferon-tau and other conceptus-originated signaling on the remodeling of the caruncle (50).

Our findings identified genes with high levels of co-expression (|r| > 0.9999) between extraembryonic tissue (n = 229) and endometrial caruncular tissues (n = 218) whose transcript profiles independently produced equivalent discrimination of the pregnancies. Functional interrogation of these 444 genes revealed that highly co-expressed genes between extraembryonic and caruncular tissues are involved in regulatory functions at the chromatin, mRNA processing, and protein levels; which is a strong indication of a coordinated reprograming of tissues driven by multiple layers of cell regulation during the conceptus-maternal recognition. These data prompt the need for additional investigation to better define the coordinated interactions between extra-embryonic tissues and endometrium at the level of tissue layer including luminal epithelium, stroma and glandular epithelium.

In the intercaruncular tissues, our analyses identified a list of genes related with “oxidation-reduction process”, a finding consistent with a recent publication reporting that proteins associated with oxidation-reduction are enriched in the uterine luminal fluid on gestation day 16 in cattle (51). Oxidative stress is a consequence of altered oxidation-reduction state (52) and transcriptional regulation of factors involved in the regulation of oxidative stress has been reported in the bovine endometrium during oestrous cycle and early pregnancy (41, 53), Furthermore, a significant increase in oxidation-reduction potential was observed in the endometrium of mice prior to implantation (54). The results show evidence that the maintenance of oxidation-reduction status permissive to the conceptus health (55) and implantation is strongly linked to genes regulated in the glandular area of the endometrium in cattle.

The analyses carried out in this study have provided novel insights into the molecular contribution of extraembryonic, caruncular, and intercaruncular tissues to conceptus elongation, uterine receptivity, and implantation, summarized in Fig 5. Gene products expressed by the extraembryonic tissue impact the endometrial function by regulating diverse cell functions including oxidative stress, chromatin remodeling, gene transcription, mRNA processing and translation. The endometrium also exerts key regulatory roles on the extraembryonic tissue cells by modulating chromatin remodeling, gene transcription, cell proliferation, translation, metabolism, and signaling (Fig 5). Collectively, our data have shown that endometrial plasticity, a notion first suggested in cattle (6), allows unique adaptive and coordinated conceptus-matched interactions at implantation in non-pathological pregnancies.

**Fig 5.**
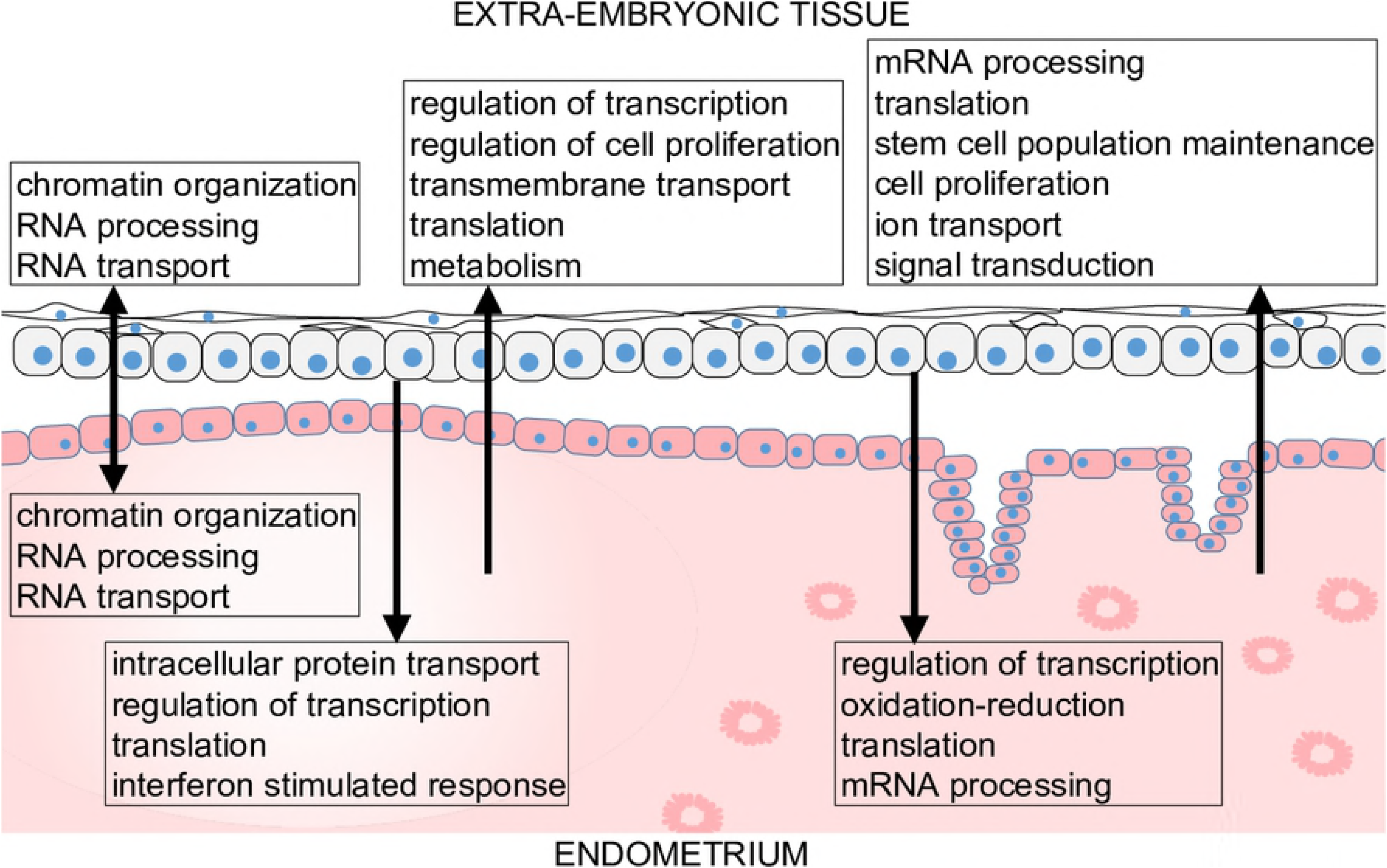
Working model of most prominent biological functions modulated by co-expression between extraembryonic tissue and endometrium. The arrows indicate probable direction of interaction.

To our knowledge, this study presents the first analysis of paired conceptus and endometrium in a mammalian species, using and integrative systems biology approach. Our results provide strong evidence that implantation in mammalian pregnancy relies on the ability of the endometrium to elicit a fine-tuned adaptive response to the conceptus in normal pregnancy. This finding opens new venues for the development of strategies to improve term pregnancy rates when artificial reproductive technologies are used. Since the endometrial response is embryo-specific, it would be valuable to develop approaches aiming at selection of the competent embryo better suitable for the establishment of a successful cross-talk with the recipient uterus of the female considered for transfer.

## MATERIAL AND METHODS

All analytical procedures were carried out in R software (36). The files and codes for full reproducibility of the results are listed on the S1 Code.

### Data analyzed and estimation of gene expression levels

The appropriated approval from institutional committees of ethical oversight for animal use in research was obtained as reported previously (26). Briefly, all five cattle gestations were initiated by artificial insemination using semen from a single bull, and later terminated on gestation day 18 for sample collection. We analyzed RNA-seq generated from samples obtained from cattle gestations interrupted at day 18 (n = 5, GSE74152). The samples were extraembryonic tissue (n = 5), caruncle (n = 5), and intercaruncle (n = 5) regions from the endometrium.

The reads were aligned to the bovine genome (*Bos taurus*, UMD 3.1) using STAR aligner (56). Reads that aligned at one location of the genome with less than four mismatches were retained for elimination of duplicates. Non-duplicated reads were used for estimation of fragments per kilobase per million reads (FPKM) using Cufflinks (v.2.2.1 (57)) and Ensembl gene models (58). Genes were retained for downstream analyses is FKPM > 1 in ≥ 4 samples. We employed the t-Distributed Stochastic Neighbor Embedding approach (59) to assess the relatedness of the tissues.

### Calculation of correlation of gene expression between tissues

Three samples were collected from the same pregnancy, thus the data structure (Fig 1B) allowed us to quantify the association between genes expressed in extraembryonic tissue and endometrium (caruncular and intercaruncular tissues). We utilized Pearson’s coefficient of correlation due to its sensitivity to outliers(60) to calculate 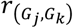 and 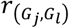, where *G_j_, G_k_*, and *G_l_*, are the transcript abundance of a gene expressed in extraembryonic tissue caruncle and intercaruncle respectively. Empirical FDR was calculated by permuting the pregnancy index (*i* = 1,…,5) for the extraembryonic tissue samples thereby breaking the pairing of conceptus and endometrium obtained per pregnancy (100 permutations) and using the formulas described elsewhere to calculate the proportion of resulting correlation resulted from the scrambled data that was greater that a specific threshold (33, 61, 62).

### Testing the resemblance of two distance matrices

We calculated distance matrices for extraembryonic tissue and caruncle passed on the Pearson’s coefficient of correlation of the expressed genes within tissues. The correlation matrix was subtracted from one to obtain a distance matrix which was used as input for clustering using the method ‘complete’. We used the Mantel statistic test implemented in the ‘mantel’ package to assess the correlation between the two dissimilarity matrices. The significance of the Mantel statistic was assessed by a permutation approach.

### Clustering of samples, heat maps, and network visualization

We clustered samples using ‘flashClust’ package (63); we used the ComplexHeatmaps package (64) to draw annotated heatmaps and Cytoscape software (65) to visualize the networks.

### Testing for enrichment of gene ontology terms or KEGG pathways

We tested for enrichment of gene ontology (66) categories and KEGG pathways (67) using the ‘goseq’ package (68). Subsets of genes were defined according to appropriate thresholds and defined as ‘test genes’; the genes expressed in the corresponding tissue were then used as background for the calculation of significance values (69). Significance values were then adjusted for FDR according to the Benjamini and Hochberg method (70).

## ACKNOWLEGEMENTS

The authors would like to thank the staff of INRA experimental units (Leudeville, France; Saint-Genes-Champanelle, France; Nouzilly, France) for animal management as well as past and present members of UMR BDR including Dr. H. Jammes, M. Guillomot, A Chaulot Talmon and Corinne Giraud-Delville for help in sample collection and preparation.

